# Implementation of an Artificial Rearing System and Molecular Identification of *Gyropsylla spegazziniana* (*Lizer & Trelles*) (*Hemiptera: Psylloidea: Aphalaridae*)

**DOI:** 10.1101/2025.07.29.667441

**Authors:** Yesica Gisel Candia, Alejandra Badaracco, Vanesa Nahirñak, María Elena Schapovaloff, Nicolás Esteban Bejerman

## Abstract

*Gyropsylla spegazziniana*, commonly known as the yerba mate psyllid, is one of the main pests affecting *Ilex paraguariensis* (yerba mate) production. To date, this pest has only been characterized morphologically, and no efficient artificial rearing system has been developed. In this study, we implemented a successful artificial rearing protocol under controlled conditions, a key step for further investigations. In addition, a fragment of the mitochondrial gene cytochrome c oxidase subunit I (COI), one of the most widely used markers for phylogenetic characterization in insects, was amplified and sequenced, enabling the first molecular identification of this pest. The obtained data confirmed the previously proposed taxonomic position of *G. spegazziniana*, which was based on morphological traits, as a member of the family *Aphalaridae*. This study lays the foundation for future research on this economically important pest species of yerba mate.

## INTRODUCTION

The superfamily *Psylloidea* (psyllids) represents an ecologically diverse but relatively understudied group within the suborder *Sternorrhyncha*, order *Hemiptera*. (Burckhardt *et al*., 2021; Mauck *et al*., 2024). Members of this group, commonly referred to as psyllids or jumping plant lice, are sap-sucking insects highly specialized in their host plants. To date, around 4,000 species have been described and classified (Burckhardt *et al*., 2021), each exhibiting distinct life histories and host exploitation strategies (Mauck *et al*., 2024). Some psyllid species are among the most devastating agricultural pests worldwide, either due to their ability to transmit plant pathogens (Bastin *et al*., 2023) or the range of phenotypes they induce in host plants through salivary secretions (Mauck *et al*., 2024).

In Argentina, around 16% of the 470 psyllid species known in the Neotropical region have been reported, and several of them are considered economically important pests (Burckhardt, 2008). *Gyropsylla spegazziniana* (Lizer & Trelles), commonly known as the yerba mate psyllid or “rulo”, is one of the major pests of yerba mate (*Ilex paraguariensis* A. St-Hil.). This crop holds substantial regional importance, as its leaves and stems are commonly used to prepare a traditional beverage known as ‘mate’ widely consumed in Argentina, Paraguay, Uruguay, and Brazil (Cardozo *et al*., 2021). *Gyropsylla spegazziniana* induces the formation of leaf deformations, referred to as “galls” or “rulos”, which serve as protective structures for its eggs and nymphs until adulthood, causing important economic losses in yerba mate production (Ohashi *et al*., 2018).

Although several studies have examined the biology, ecology, morphology, and control of *G. spegazziniana*, (Sabedot *et al*., 2000; Leite & Zanol, 2001; Leite *et al*., 2007; Alves *et al*., 2009, 2013, Formentini *et al*., 2015; Ohashi *et al*., 2018; Queiroz *et al*., 2021), molecular data and phylogenetic relationships remain lacking. Furthermore, while some laboratory protocols have been tested under standard conditions for short periods (Formentini *et al*., 2015; Loeblein *et al*., 2019), no efficient artificial rearing system has been reported. According to Viscarret & López (2020), establishing insect colonies under controlled conditions is essential for research on pest management and control, as well as for studies on insect biology and behaviour.

In parallel, the application of molecular tools has revolutionized insect systematics. Mitochondrial markers, especially the cytochrome c oxidase subunit I (COI) gene, have proven to be powerful for phylogenetic and population genetics analyses due to their maternal inheritance, lack of recombination, and high mutation rate (Hebert *et al*., 2003). This gene, targeted by universal primers (Folmer *et al*., 1994), is the standard marker in DNA barcoding approaches, enabling precise species identification, even in morphologically conserved or cryptic lineages such as psyllids (Pramatarova *et al*., 2024).

Therefore, the present study aimed to implement a continuous artificial rearing system for *G. spegazziniana* under controlled conditions and to perform molecular identification using COI-based phylogenetic analysis, providing insights into the systematic placement of this species within *Psylloidea*.

## MATERIALS AND METHODS

### Artificial Rearing System for *Gyropsylla spegazziniana*

#### Host plant cultivation

Yerba mate seedlings were obtained from nurseries in Misiones Province, Argentina, and transplanted individually into 1-L transparent pots containing soil substrate and slow-release fertilizer. Plants were watered with 20 ml of tap water every day and maintained under greenhouse conditions.

#### Establishment of a founding colony under controlled conditions

As a first step, galls were field collected from yerba mate plant branches (*Ilex paraguariensis*) (Fig 1a) and placed in polypropylene bags for transport to the laboratory. Following the procedure described by Loeblein et al. (2019), the galls were partially opened manually, and those containing IV–V instar nymphs were placed into plastic containers (5 cm height × 6 cm diameter) with mesh lids and a piece of absorbent paper at the base (Fig 1b). The containers were maintained in a climate-controlled room at 26 ± 1□°C, with a 12:12 hours light:dark photoperiod and 60 ± 10% relative humidity, for approximately 24 hours until adult emergence (Fig. 1c and 1d). Subsequently, the emerged adults were transferred to a rearing cage (50 × 60 × 60 cm) constructed with anti-aphid mesh and containing yerba mate seedlings, establishing the first founding colony (“colony 1”). After completing the life cycle (± 30 days), new galls induced by these adults were collected, and the procedure was repeated to establish “colony 2,” maintained under the same controlled conditions.

**Figure 1.**
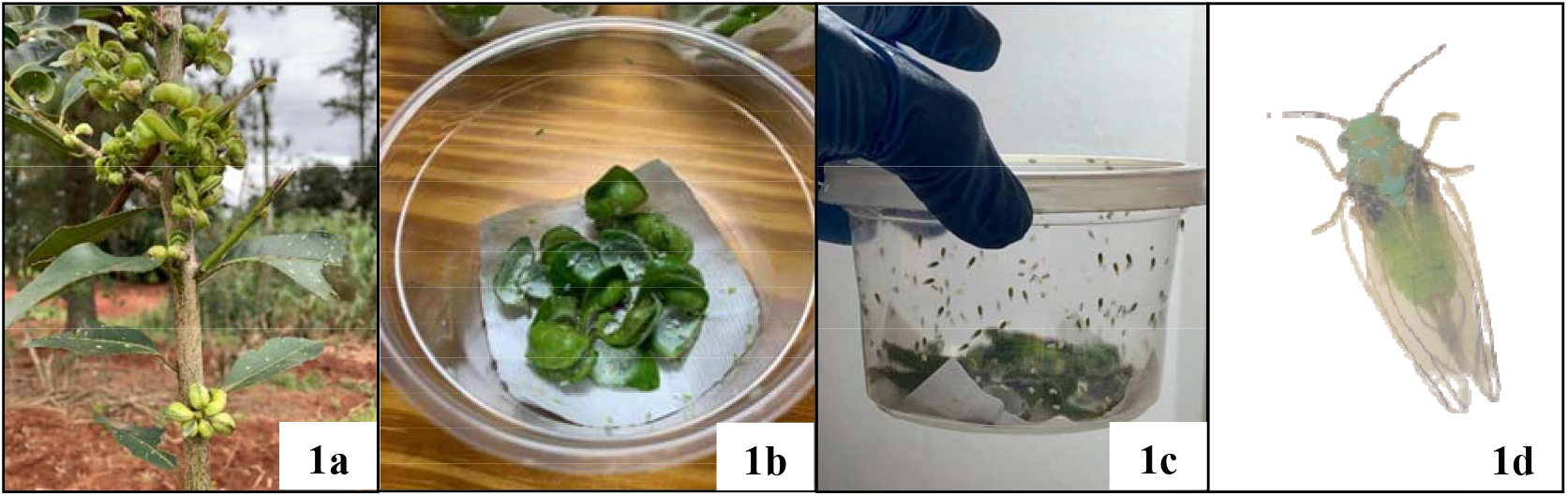
Establishment of the initial rearing colony of Gyropsylla spegazziniana under controlled conditions. 1a. Galls on yerba mate branches. 1b. Galls placed in plastic containers.1c. G. spegazziniana adults emerged after 24 hours. 1d. Adult individual of G. spegazziniana.

#### Establishment of the rearing colony under greenhouse conditions

A rearing cage was prepared using aluminium frames and sliding glass doors, with anti-aphid mesh walls (4.8 m × 1.2 m × 1 m), and equipped with yerba mate plants in 3 L pots. Adults that emerged from the galls of “colony 2” were transferred to this cage, establishing the third rearing colony, which was maintained under greenhouse conditions.

#### DNA extraction, PCR amplification, and sequencing

Genomic DNA was extracted from a single *G. spegazziniana* individual using a modified CTAB protocol (Doyle & Doyle, 1987). A fragment of the COI gene was amplified following the protocol by Folmer *et al*. (1994), using universal primers: LCO1490: 5^′^-GGTCAACAAATCATAAAGATATTGG-3^′^ and HCO2198: 5^′^-TAAACTTCAGGGTGACCAAAAAATCA-3^′^.

PCR products were visualized via 1.5% agarose gel electrophoresis in 0.5X TBE buffer stained with GelRed™ (Biotium, 10,000X). Amplified fragments were purified using the EasyPure Quick Gel Extraction Kit (TRANS®), following the manufacturer’s instructions. Sanger sequencing was performed by Macrogen Inc. (Seoul, South Korea; http://macrogen.com).

#### Sequence database construction and phylogenetic analysis

The obtained sequences were compared against those in the GenBank database (NCBI; https://www.ncbi.nlm.nih.gov) using the BLASTn algorithm (Altschul *et al*., 1990). Multiple sequence alignments were conducted using MUSCLE in MEGA11 software (Tamura *et al*., 2021). A phylogenetic tree was built using the Neighbor-Joining method with 1,000 bootstrap replicates to assess branch support. Evolutionary distances were estimated using the JTT+G substitution model.

## RESULTS AND DISCUSSION

### Rearing system

Under controlled conditions, colonies 1 and 2 were successfully established (Fig. 2), while the third colony was obtained under greenhouse conditions (Fig. 3a). In all rearing colonies, a continuous flow of *G. spegazziniana* individuals was observed at 28 - 35 day intervals, with the presence of induced galls (Fig. 3b) and live adults in each generation (Fig. 3c). These results demonstrate that the adults obtained through rearing were reproductively functional and that the designed environment was suitable for their maintenance. Although survival and fecundity were not exhaustively quantified, consecutive days with active individuals in the cages were recorded, consistent with the findings of Sabedot *et al*. (2000), who reported a life cycle of approximately 28 days. Leite & Zanol (2001) observed a slightly longer cycle (38 days), a difference that may be attributed, as noted by the authors, to manipulation methods and host plant availability.

**Figure 2.**
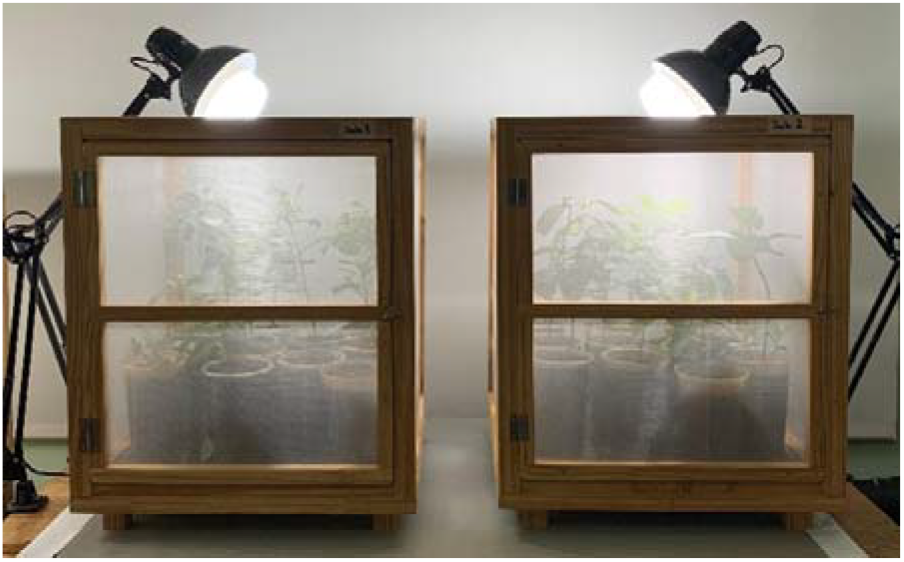
Colonies 1 and 2 maintained under controlled environmental conditions.

**Figure 3.**
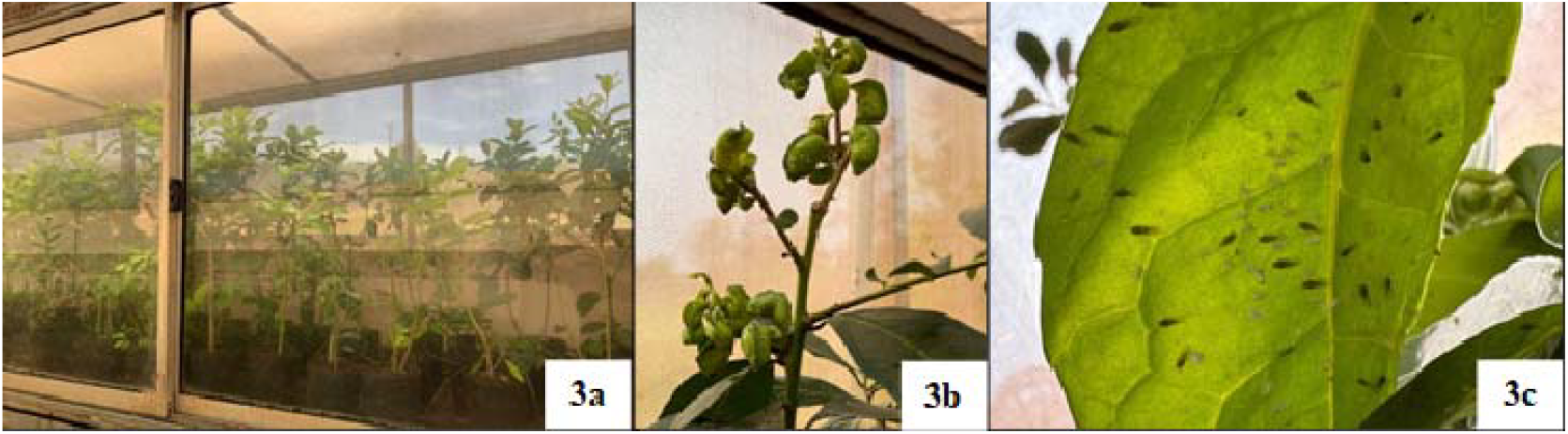
Colony 3 maintained under greenhouse conditions. 3a. Rearing cage containing Ilex paraguariensis seedlings.3b. Galls formed on yerba mate plants inside the rearing cage.3c. G. spegazziniana adults posed on a yerba mate leaf.

Although the procedure in this study was based on the previously described by Loeblein *et al*. (2019), it is important to emphasize that this study represents the first successful attempt at establishing a complete artificial rearing system for *G. spegazziniana*. Loeblein *et al*. (2019) focused only on gall manipulation and adult emergence under laboratory conditions, without establishing multi-generational colonies. The continuity achieved here confirms the feasibility of the proposed protocol and its potential utility for future research.

Artificial rearing systems have also been successfully implemented for other insect species. For example, *Diaphorina citri* and *Tamarixia radiata* were reared under controlled conditions for behavioural assays related to biological control strategies (Skelley & Hoy, 2004; Aguirre, 2019). These studies highlight the importance of maintaining stable colonies for both basic and applied research.

A reliable artificial rearing system is an essential requirement for implementing control strategies (biological, genetic, or chemical), as well as for studies in insect biology, virology, ecology, and reproduction (Viscarret & López, 2020). The ability to maintain individuals year-round reduces reliance on field collection and facilitates repeatable and controlled experiments.

### Molecular identification

A fragment of the cytochrome c oxidase subunit I (COI) gene was successfully amplified in *G. spegazziniana* using the universal primers LCO1490 and HCO2198 (Folmer *et al*., 1994). The amplified product was 708 base pairs in length, and the sequence was deposited in GenBank under the accession number PV925725.

BLAST analysis of the amplified fragment showed the best match with the COI gene of *Aphalara* sp., with 86.13% identity. The phylogenetic tree constructed using the Neighbor-Joining method supported the BLAST findings, placing *G. spegazziniana* within the clade corresponding to the family *Aphalaridae*, grouped with species from the genera *Aphalara, Craspedolepta*, and *Lanthanaphalara* (Fig. 4). This strongly supports its classification within the family *Aphalaridae*. The grouping suggests a close phylogenetic relationship with members of this family, consistent with previous classifications based on morphological traits (Burckhardt *et al*., 2021) and highlights the value of the COI gene as a reference marker for accurate taxonomic identification of psyllids (Pramatarova *et al*., 2024). Our study constitutes the first molecular report for this species, reinforcing its phylogenetic placement and providing a genetic foundation for future investigations.

**Figure 4.**
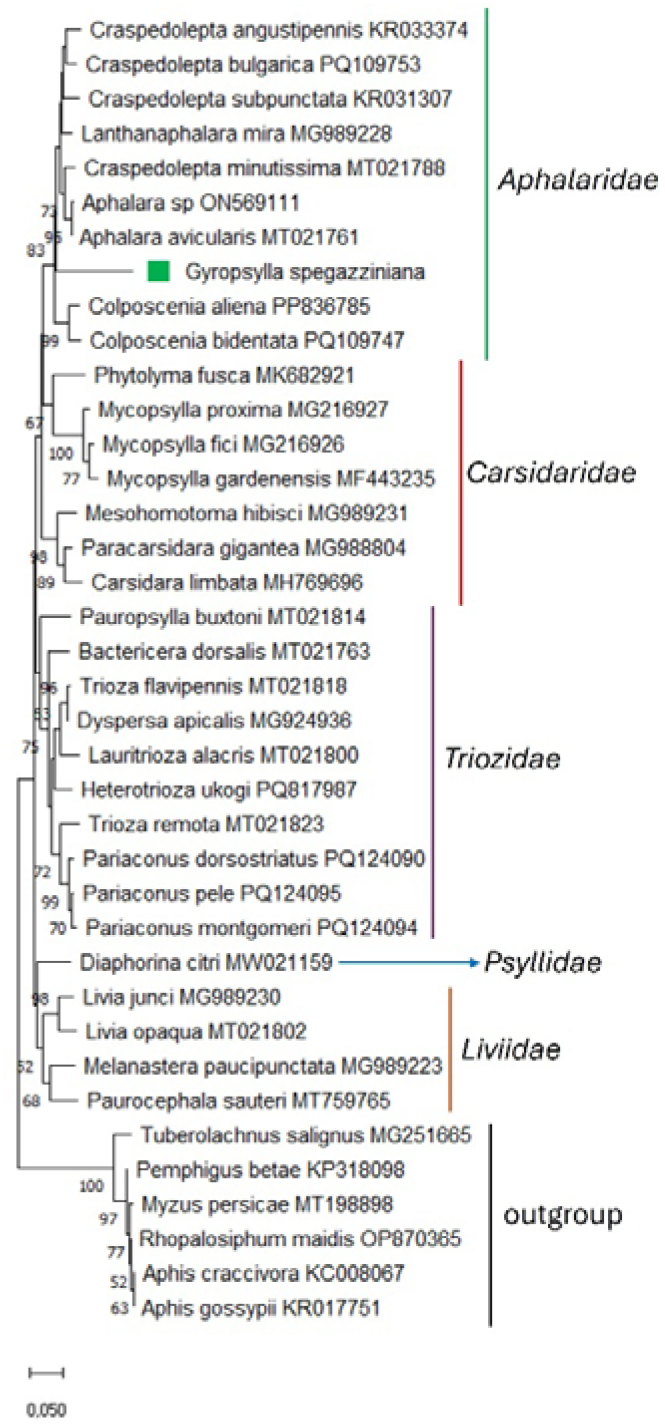
Phylogenetic tree using the Neighbour-joining method.

The results presented here represent a significant advance, as the establishment of an artificial rearing system for *G. spegazziniana* offers a valuable tool for future studies on this economically important pest. Additionally, given the absence of prior molecular data for this psyllid, the successful amplification of the COI gene offers a reliable marker for its molecular identification. This significantly broadens the scope for future research into its biology and the development of sustainable management strategies for yerba mate cultivation.

## ACKOWLEDGEMENTS

This project was funded by the Instituto Nacional de Yerba Mate (INYM) through the PRASY project “Sequencing and analyzing the transcriptome of the yerba mate pest *Gyropsylla spegazziniana*”.

## DATA AVAILABILITY STATEMENT

The generated COI data is available in the GenBank of the NCBI under accession number PV925725.1.

